# A spatially resolved stochastic model reveals the role of supercoiling in transcription regulation

**DOI:** 10.1101/2021.12.29.474406

**Authors:** Yuncong Geng, Christopher H. Bohrer, Nicolás Yehya, Hunter Hendrix, Lior Shachaf, Jian Liu, Jie Xiao, Elijah Roberts

## Abstract

In *Escherichia coli*, translocation of RNA polymerase (RNAP) during transcription introduces supercoiling to DNA, which influences the initiation and elongation behaviors of RNAP. To quantify the role of supercoiling in transcription regulation, we develop a spatially resolved supercoiling model of transcription, describing RNAP-supercoiling interactions, topoisomerase activities, stochastic topological domain formation, and supercoiling diffusion in all transcription stages. This model establishes that transcription-induced supercoiling mediates the cooperation of co-transcribing RNAP molecules in highly expressed genes. It reveals that supercoiling transmits RNAP-accessible information through DNA and enables different RNAP molecules to communicate within and between genes. It thus predicts that a topological domain could serve as a transcription regulator, generating substantial transcription bursting and coordinating communications between adjacent genes in the domain. The model provides a quantitative platform for further theoretical and experimental investigations of how genome organization impacts transcription.

**Author Summary:** DNA mechanics and transcription dynamics are intimately coupled. During transcription, the translocation of RNA polymerase overwinds the DNA ahead and underwinds the DNA behind, rendering the DNA supercoiled. The supercoiled DNA could, in return, influences the behavior of the RNA polymerase, and consequently the amount of mRNA product it makes. Furthermore, supercoils could propagate on the DNA over thousands of base pairs, impacting RNA polymerase molecules at faraway sites. These complicated interplays between supercoiling and RNA polymerase makes supercoiling an important transcription regulator. To quantitatively investigate the role of supercoiling in transcription, we build a spatially resolved model that links transcription with the generation, propagation, and dissipation of supercoiling. Our model reveals that supercoiling mediates transcription at multiple length scales. At a single-gene scale, we show that supercoiling gives rise to the collective motion of co-transcribing RNA polymerase molecules, supporting recent experimental observations. Additionally, large variations in mRNA production of a gene can arise from the constraints of supercoiling diffusion in a topological domain. At a multi-gene scale, we show that supercoiling dynamics allow two adjacent genes influence each other’s transcription kinetics, thus serving as a transcription regulator.

## Introduction

Recent studies show that in *E. coli* the chromosomal DNA is under torsional stress. A relaxed DNA (with every 10.5 bp per turn [1]) becomes torsionally stressed if it is underwound (negatively supercoiled) or overwound (positively supercoiled). In cells, the chromosomal DNA is held in a homeostatic, negatively supercoiled state globally by the coordination of topoisomerases [2, 3]. However, DNA-related processes such as replication and transcription could drive the local supercoiling state away from its equilibrium. For example, the twin-domain model of transcription depicts that during elongation, RNA polymerase (RNAP) overwinds the DNA downstream and underwinds the DNA upstream to generate positive and negative supercoiling respectively, leading to the build-up of DNA torsional stresses in the vicinity of RNAP [4, 5]. During this process, RNAP “encodes” torsional information on the DNA. If the torsional stress is not relieved in a timely manner, it could propagate through the DNA and influence the behavior of other DNA-bound RNAP molecules from afar [6]. RNAP “retrieves” the torsional information by adapting its initiation and elongation behaviors in response to local supercoiling. Many promoters display supercoiling sensitivity [7, 8, 9], and high torsional stress could slow down [8] or even stall [10, 11] elongating RNAP molecules.

The unique property of supercoiling to store and transmit RNAP-accessible information makes it a medium for RNAP molecules on the same transcription unit or even different operons kilobases away to communicate with each other [12, 13]. Therefore, supercoiling has been proposed to serve as a transcription regulator across multiple distance scales from a few to thousands of base pairs, giving rise to many emergent properties in transcription.

At the single-gene scale, supercoiling is proposed to coordinate the collective motion of RNAP molecules during elongation, enabling RNAP to move at different speeds with different initiation rates. The promoter strength, in this case, emerges as a regulator for the elongation speed. Kim *et al.* [13] found that strong promoters facilitate the processivity of RNAP when the promoter is on and inhibits the processivity of RNAP when the promoter is off. The authors hypothesized that during transcription, opposite supercoils generated in the region between two adjacent RNAP molecules can cancel out with each other quickly, reducing the torsional stress in the region and hence making the translocation of both RNAP molecules faster.

At the multi-gene scale, supercoiling could mediate the interaction between multiple closely arranged genes. Genome-wide evidence of correlation of transcriptional activity between neighboring genes has been detected [14, 15], and the transcription of gene cassettes inserted in the *E. coli* genome was found to depend strongly on the transcription activity of adjacent genes at the insertion site [16, 17]. It is suggested that supercoiling plays a role in producing the correlation — transcribing RNAP molecules of adjacent genes keep rewriting the intergenic supercoiling profile and preventing their dissipation, hence enabling neighboring genes to “feel” and affect the transcription of each other through the sharing of the same torsional stress state [14, 15, 16].

Supercoiling could also regulate the transcription of genes belonging to the same topological domains. In *E. coli* cells, there are about 400 topological domains of an average size of ~10kb [18, 19], which are likely formed by the dynamic looping and unlooping of the intervening DNA due to the binding and unbinding of nucleoid-associated proteins. Those proteins, by the nature of their binding to the DNA, restrict the diffusion of supercoiling and hence constrain torsional stress within topological domains. Supporting this possibility, theoretical work has shown that the opening and closing of a topological domain could lead to transcriptional bursting [20] and correlate gene transcription in the same domain [21].

Although supercoiling is proposed to mediate the communication between multiple RNAP molecules and multiple genes, it is not clear how supercoiling dynamics lead to changes in transcription activities quantitatively and how these changes are dependent on the mechanical properties of DNA and the transcription kinetics of RNAP. Several models [20, 21, 22, 23, 24, 25, 26, 27] have been established to explore the role of supercoiling in transcription regulation. However, none of these models fully captured the complex interactions between transcription and supercoiling. Instead, they mainly focused on a subset of key aspects, such as the contribution of supercoiling to transcription initiation [21, 23, 24, 26] or elongation [20, 22, 25, 27] only. Here we present a spatially resolved, chemical master-equation based model of *E. coli* transcription that explicitly describes all stages of transcription (initiation, elongation, and termination, **Fig 1A-C**), and their interplay with DNA supercoiling and other cellular processes that regulate DNA torsional stress (topoisomerase activity, supercoiling diffusion, and the dynamical formation and dissociation of topological domains, **Fig 1D-G**). This comprehensive model integrates the most recent experimental observations of the relationships between transcription and supercoiling.

**Fig 1.**
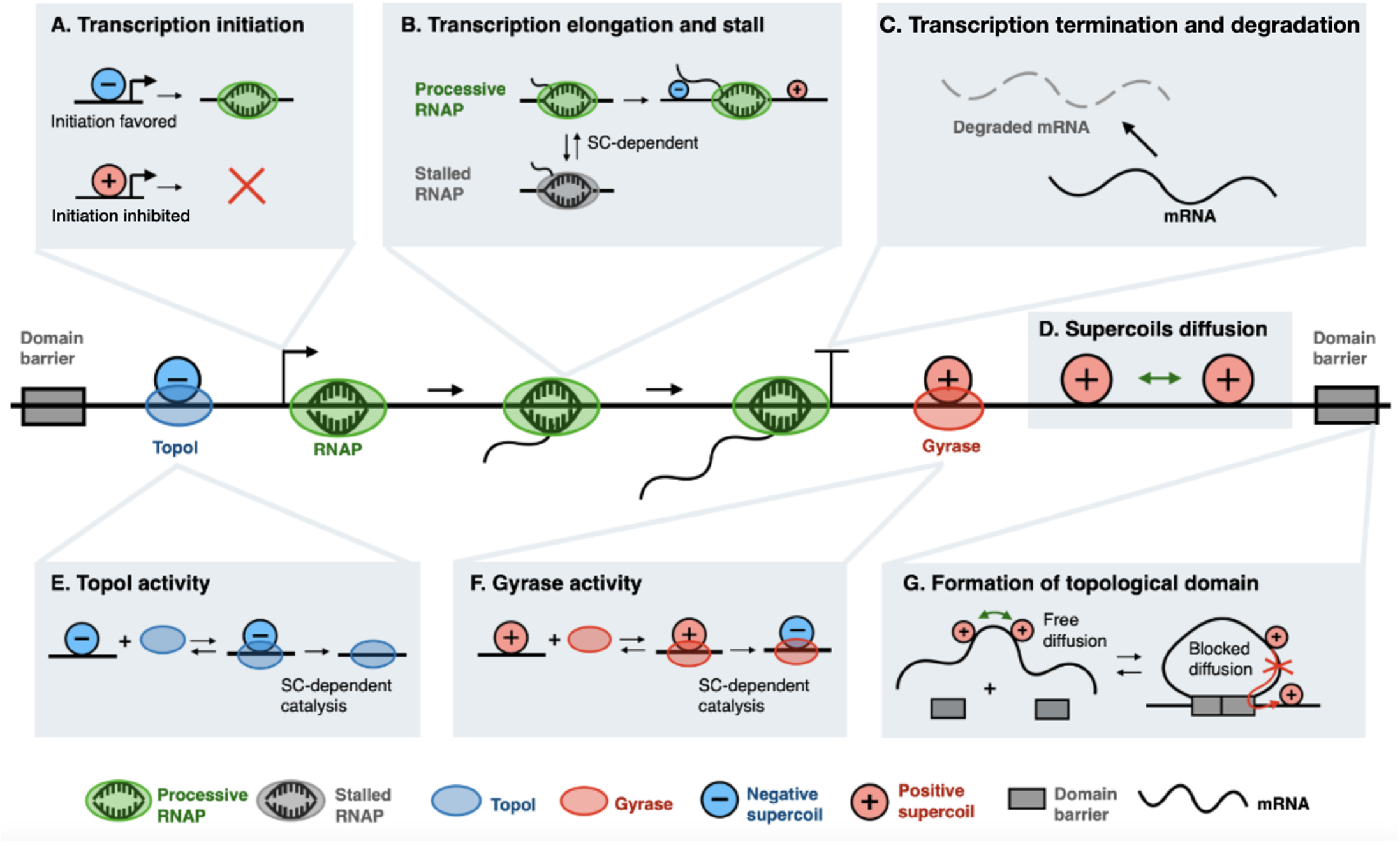
Model schematics of supercoiling-dependent transcription. (A) During transcription initiation, a negatively supercoiled (blue circle) promoter is favored over a positively supercoiled (red circle) promoter for RNAP (green oval) binding and open complex formation. (B) During transcription elongation, the translocation of RNAP induces positive supercoiling in front and negative supercoiling behind, which consequently influence whether the RNAP remains processive (green) in elongation or becomes stalled (grey). (C) Transcription terminates when RNAP is released from the mRNA. The transcribed mRNA is degraded subsequently in the model. (D) Supercoils can diffuse on DNA within a topological domain flanked by two domain barriers and interact with RNAP and topoisomerases. (E) Topo I (blue oval) removes one negative supercoil at a time. (F) Gyrase (red oval) converts one positive supercoil to one negative supercoil at a time. (G) Formation and dissolution of a topological domain upon the binding and unbinding of domain anchoring proteins (domain barriers, grey block).

Using this model, we quantitatively established that inter-RNAP supercoiling dynamics could fully account for the cooperation of co-transcribing RNAP molecules. We also predicted that by confining supercoiling diffusion, a topological domain can give rise to bursty transcription of genes within the domain. Furthermore, transcription-induced supercoiling leads to complex communications between two closely arranged genes. Our findings provide insights into how supercoiling regulates gene expression in the context of promoter architecture and gene arrangement, paving the way to quantitative studies of how genome organization impact gene expression.

## Methods

### General design of the model

The model describes the mechanochemical feedback between RNAP activities and DNA supercoiling, where supercoiling controls the chemical reaction rates of RNAP, which in turn modulates the mechanics of DNA and hence supercoiling. As both the supercoiling and the RNAP molecules diffuse laterally, such mechanochemical feedback plays out along the DNA. The model thus evaluates how such feedback modulates and coordinates transcriptions along the DNA. Below we provide the mathematical formulation of the model.

We implemented a modified reaction-diffusion model using chemical master equations, treating the DNA as a linear 1-D lattice. Specifically, we divided the DNA into a series of 60-bp segments; the choice of 60-bp is to a compromise to accommodate the length scale of both RNAP footprint (32 bp) and gyrase binding site (137 bp) and to minimize computational costs. Supercoils, RNAP or topoisomerase molecules at different positions of the lattice were considered as different species. This strategy enabled us to describe the spatial distribution of RNAP, topoisomerases, and supercoils at different positions along the DNA over time. We performed simulations with exact sampling (Gillespie algorithm) in Lattice Microbes [28], with counts of species recorded every 1 s. Due to the limited number of species allowed in Lattice Microbes, the maximum size of DNA we simulated was at ~ 20 kb. The detailed rationale of this strategy can be found in **S1 Text**.

To describe the dynamics in the coupling of transcription and supercoiling, we focused on two questions: (1) how DNA supercoils are generated and dissipated by transcription and other processes, and (2) how transcription responds to DNA supercoiling. To address the first question, we incorporated into our model three processes that are known to contribute to the dynamics of DNA supercoiling: (a) diffusion of supercoils, (b) activities of topoisomerases, and (c) transcription. Topoisomerases regulate supercoiling globally, transcription perturbs supercoiling locally, and through diffusion, those changes are transmitted along the DNA. To address the second question, our model considered the supercoiling-sensitive transcription initiation and elongation. Detailed reaction schemes are described below. All model equations and parameters can be found in **S1 Table**, **S2 Table** and **S3 Table**.

### Modeling diffusion of supercoiling

DNA torsional stress can be stored in the form of either twist or writhe [29]. Twist is the rotation of DNA around its rise axis. Writhe is the cross-over of DNA double-helix with each other. When a sufficiently high torque is applied to DNA, instead of twisting, DNA will wrap around itself to form writhes, since the energetic cost of bending becomes lower than that of twisting [30] (**S1 Fig**). Further increases in the torque will cause the writhes to pile up and form plectonemes [32]. Twists and writhes diffuse differently along DNA. Twists diffuse rapidly along the DNA, with an estimated diffusion constant of 50 ~ 180 *μm^2^* · *s*^−1^ [34, 35]. Writhes diffuse slowly, with a very small diffusion coefficient *(D)* of 0.01 *~* 0.2 *μm*^2^ · *s*^−1^, according to *in vitro* experiments performed by Loenhout *et al.* [33]. Because DNA exists mostly in the form of writhes (or plectonemes) under physiological conditions [36], we assume that torsional stress diffuses relatively slowly.

In our model, we used “supercoiling density” (*σ*) to quantify the level of torsional stress, and we used the universal term “turns” to represent both twists and writhes. Supercoiling density is defined as 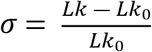, where *Lk* is the actual number of turns in the DNA segment, and *Lk_o_* is the number of turns in the DNA when it is fully relaxed (one turn per 10.5 bp). A relaxed, 60 base pair DNA segment has a *Lk_o_* at 60 bp/(10.5 bp/turn) = 5.7 turns/segment. In the *E. coli* chromosome, the average supercoiling density *σ* is at −0.05 ~ −0.07 [3], corresponding to a *Lk* at ~ 5.4 turns/segment.

We used DNA(*k*) to denote the availability of the *k*-th DNA segment. DNA(*k*)=1 indicates that the DNA is unoccupied and available for the binding of RNAP, Gyrase or topoisomerase, and DNA(*k*)=0 indicates that the DNA segment is occupied by a DNA binding-protein and unavailable. We used Turn(*k*) to track the total number of turns on the *k*-th DNA segment. To reduce the computational load, we modeled the diffusion of turns as a biased random walk, which means that we only allowed turns to move down the gradients:

If Turn(*k*)>Turn(*k-1*) and DNA(*k-1*)=1 (unoccupied), the turns on the *k-th* segment diffuse to (*k-1*)-th segment:

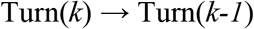

Similarly, if Turn(*k*)>Turn(*k+1*) and DNA(*k+1*)=1:

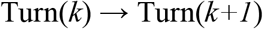

The rate at which turns diffuse along the DNA segments is determined by the propensity of the above two reactions and the rate constant for turn displacement, *k_drift_*. We set *k_drift_* as 50 *s*^−1^, which yields a diffusion coefficient *D* of 0.02 *μm^2^* · *s*^−1^, in accordance with previously estimated values [33].

### Parameterizing the binding kinetics and catalytic activities of topoisomerases

We decomposed topoisomerase activities into three steps: binding to DNA (with the rate constant of *k_bind_)*, catalysis (with the rate constant of *k_cat_)*, and dissociation (with the rate constant of *k_unbind_)*. Below we deduced the values of the three rate constants from the existing data. We assumed that topoisomerase binds indiscriminately throughout DNA, and only the catalysis rate depends on the local supercoiling density. Stracy *et al*. [37] recently characterized the binding kinetics of Gyrase in *E. coli* using single-particle tracking. There are about 300 Gyrase molecules stably bound to the DNA at any time, and the average dwell time is about 2 s. Since the *E. coli* genome is about 4 Mb, taking a deterministic approximation, we have *k_unbind_* = 1/(2 s) = 0.5 *s*^−1^, and *k_bind_* = (300/(4 Mb/(60 bp/segment)))/(2 s) = 0.00225 *s^−1^·segment*^−1^. Because of the deterministic approximation, we further corrected *k_bind_* to 0.0018 *s^−1^·segment^−1^* to match published experimental results [37]. For TopoI’s binding kinetics, no experimental measurements are available, therefore we approximated the unbinding rate of TopoI as the same of Gyrase, and treated the binding rate as a free parameter. The response of catalysis activity to supercoiling density *k_cat_(σ)* for both TopI and Gyrase was parameterized from the DNA relaxation assays (**S1 File**) [8, 38].

### Modeling the effects of transcription on supercoiling

In *E. coli*, a transcribing RNAP molecule usually forms a macromolecular complex with a nascent mRNA molecule, translating ribosomes, and newly translated peptides. As such, a transcribing RNAP molecule is, in general, considered as a topological barrier that blocks the diffusion of supercoiling. A transcribing RNAP molecule can travel linearly along the DNA to generate torsional stress, and/or counterrotate to release torsional stress. Our model considers both situations as described below.

Assuming that an RNAP molecule does not rotate at all, the linking number upstream and downstream of it would be conserved. As a result, when an RNAP molecule travels along the DNA, it will push all the turns downstream forward instantaneously and introduce torsional stress (**S2 Fig**). Since RNAP displacement is powered by the chemical reactions of NTP’s incorporation, and the concentration of NTP is held constant in most cases, we expected a constant translocation rate (60 *bp·s^−1^*). To avoid collisions between multiple RNAP molecules and the turns of each DNA segment pushed by RNAP, we only allowed displacement of RNAP molecules to occur when the two downstream segments (namely, DNA(k+1) and DNA(k+2)) are not occupied by other RNAP molecules.

In addition to directly displacing turns on adjacent DNA segments, RNAP could also counterrotate itself to release torsional stress [8]. In the *in-vitro* study by Chong *et al*. [8], it was observed that the transcription initiation rate of a circular DNA remained constant in the absence of topoisomerase even when both ends were fixed to a surface to avoid free rotations. In this case, as an RNAP molecule approaches the end of a gene that is fixed to the surface by biotin tags (serving as topological barriers), an extreme level of positive supercoiling would build to the extent that stalls transcription elongation and subsequently initiation if RNAP does not counterrotate to release the stress. Premature dissociation of RNAP could also be one way to release the torsional stress, but it is unlikely because it is well known that the transcription elongation complex is extremely stable and unlikely to dissociate spontaneously without the help from active release factors *in vivo* [39]. Here we modeled the counterrotation of RNAP using the leaky diffusion of supercoils over RNAP, and we defined the rotation rate *k_rot_* as the leaky diffusion rate (**S2 Fig**). Since we know very little about the rotation rate of RNAP, we kept it as a free parameter. We also assume that RNAP’s displacement and rotation are uncoupled.

### Modeling supercoiling-sensitive transcription initiation

Open complex formation is a crucial step in transcription initiation, and promoter-melting is favored by negatively supercoiled DNA [7]. Recently, using *in vitro* single-molecule fluorescence *in-situ* hybridization (smFISH), Chong *et al*. [8] confirmed that both the T7 RNAP and the *E. coli* RNAP have supercoiling-sensitive initiation rates. These experiments motivated theoretical studies to quantify the influence of supercoiling on transcription initiation. By adopting a statistical mechanics model, Bohrer *et al*. [21] approximated the initiation rate *k_i_*(*σ*) by a linear function of the supercoiling density *σ* at the promoter. This model quantitatively recapitulated the dynamics of initiation rate observed by Chong *et al*. [8]. We leverage this formulation here to describe the supercoiling-sensitive initiation:

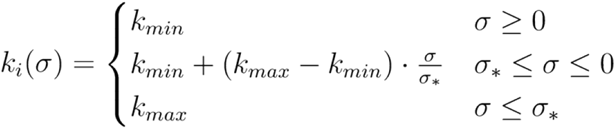

Where *σ*\* is a critical level of negative supercoiling density at which the initiation rate reaches its maximum. The initiation rate reaches its minimum when DNA is fully relaxed or further positively supercoiled. In the following simulations, *σ*\* is held at −0.06.

### Modeling supercoiling-sensitive transcription elongation

During transcription, the change in downstream and upstream DNA supercoiling generated by RNAP translocation will introduce torque around the RNAP. When the torque is beyond a certain threshold, RNAP will stall instantaneously. After the torque is released, transcription will resume. Ma *et al*. [10, 11] has shown that *E. coli* RNAPs can endure torques up to about 10.5 *pN* · *nm*. We used this value as the stall torque threshold. In our model, the torque that an RNAP molecule faces was calculated from the supercoiling density, and the details can be found in **S1 File**.

Besides torque, we also assumed that an elongating RNAP molecule could stall under other circumstances caused by extreme supercoiling densities. For example, when the upstream DNA is hyper-negatively supercoiled, R-loop might form between the template strand and the nascent mRNA [40], and hence arrest RNAP [41]. Additionally, when downstream DNA is hyper-positively supercoiled, the free energy required to melt DNA could be extremely high for RNAP to proceed [8]. To account for these effects, we set σ = ± 0.6 as the supercoiling density threshold that leads to stalled RNAP.

## Results

### RNAP Translocation-induced supercoiling mediates the collective behaviors of co-transcribing RNAP molecules

Kim *et al*. [13] found that ‘‘communication” between co-transcribing RNAP molecules on highly expressed genes manifests in two forms: cooperation and antagonism. Cooperation is the phenomenon that when the promoter is on (i.e., there is a continuous transcription initiation), each transcribing RNAP molecule elongates faster than the case when it moves solo. Antagonism is the phenomenon that when the promoter is off (i.e., no new transcription initiation), existing transcribing RNAP molecules elongate significantly slower than the case when the promoter is on. Kim *et. al*. proposed that these phenomena were due to the presence or absence of continuous loading of RNAP molecules onto the genes: the upstream negative supercoils produced by the last RNAP molecule could be rapidly canceled out by the downstream positive supercoils generated by a newly loaded RNAP molecule if the promoter is on, reducing the torsional stress, but remain unresolved if the promoter is turned off.

To explore whether the dynamics of transcription-induced supercoiling could indeed account fully for those collective behaviors, we simulated the transcription of *lacZYA* (the same systems used in Kim *et al*. [13]) centered in a 9 kb-long open domain, with both ends relaxing to the equilibrium supercoiling density - 0.067 [3] rapidly (every 0.02 s). The rotation rate of RNAP was set at 0.2 *s*^−1^, and the Topo I binding rate as 10 times of the Gyrase binding rate. To represent promoters with different strengths, we varied the maximum initiation rate constant *k_max_* from 0.001. *s*^−1^ to 0.2 *s*^−1^, which is within the strength range of *E. coli* promoters [42]. The minimum initiation rate constant *k_min_* was held as 0.001 *s*^−1^ for all simulations. The elongation rate of a group of RNAP molecules is defined as the average elongation rate of this RNAP group.

**Fig 2A** shows that genes under higher initiation rates tend to have high elongation rates as well, recapitulating the cooperative behavior of RNAP molecules. Importantly, the cooperation is non-additive, i.e., the elongation rate ‘‘saturates” when the initiation rate is sufficiently high, which agrees with Kim *et al.’s* observations [13]. Consistent with this observation, the correlation in the displacement of adjacent RNAP molecules increased and plateaued at high initiation rates (**Fig 2B**, **Supplementary Notes, Fig. SX**), indicating that RNAP tends to adjust its speed to mimic the speed of its neighbors, i.e., “cooperating” with each other. The distance between adjacent RNAP molecules continued to decrease in response to increased initiation rate (**Fig 2C**). Interestingly, the average distance between two adjacent RNAP molecules was around 500 bp when the elongation rate reached the plateau and continued to decrease further in response to increased initiation rates, indicating that the full cooperation between adjacent RNAP molecules could operate over a distance of > 500 bp.

**Fig 2.**
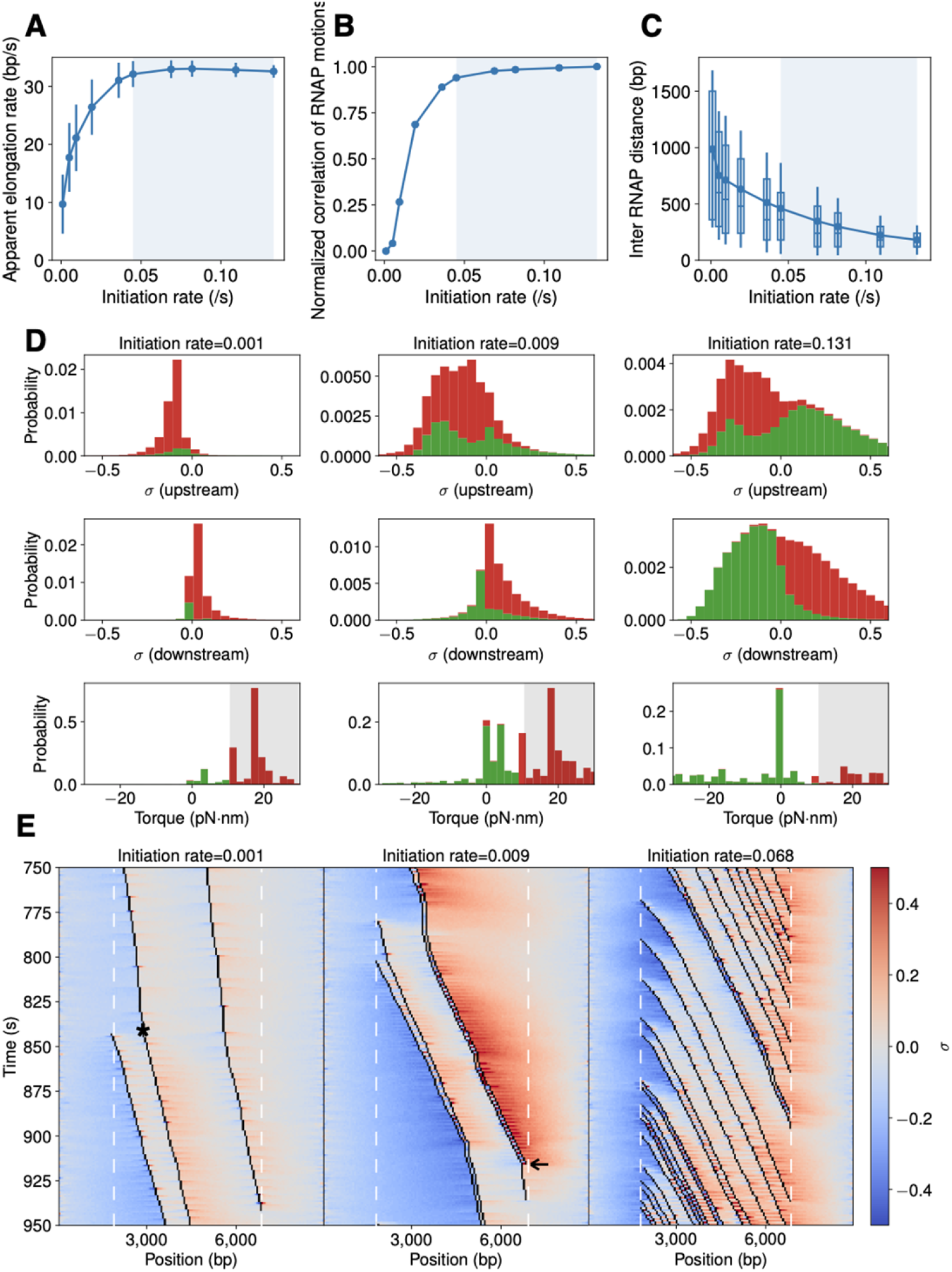
Cooperation of RNAP transcription is favored by high initiation rates and antagonized by low initiation rates. (A) The apparent RNAP elongation rate increases and reaches maxima (blue shaded region) as the initiation rate increases. Error bars are standard deviations from (x simulations?). (B) Pearson correlation coefficient of adjacent RNAP trajectories increases and approaches unity (blue shaded region) as the initiation rate increases. The correlation coefficient is normalized such that “0” corresponds to the case of independently transcribing RNAP trajectories and “1” corresponds to the case of the co-transcribing RNAP trajectories in the highest expressed gene. The unnormalized curve is in **S4 Fig**. (C) The mean distance (dots in boxes) between neighboring RNAP molecules decreases as the initiation rate increases. When the maximal cooperation is reached (intersect with blue shaded region, initiate rate at which maximal elongation rate is reached) the distance is ~ 500 bp. Box borders indicate first and third quartiles of the data, vertical lines standard deviations, and horizontal lines in the box media. (D) Stacked histograms of supercoil density upstream (top), downstream (middle) and torque (bottom) RNAP molecules face during transcription. Supercoil densities of processive and stalled RNAP molecules are shown in green and red, respectively. The torque value above the stall threshold is shaded in grey. (E) Representative kymographs of RNAP translocation trajectories (black lines) with the corresponding supercoiling density (positive in orange and negative in blue) in its vicinity. A time interval of 750 – 950 s was chosen to maintain a constant mean mRNA copy number at a steady-state under each condition (**S5 Fig**). The slope of the trajectory reflects the elongation speed of the associated RNAP molecule. Left and right white dashed lines indicate the start and end positions of the transcribing region, respectively. The asteroid in the left kymograph indicates that a slow-moving RNAP molecule started to move with a higher speed after a new RNAP molecule is loaded at the promoter on the left. The arrowhead in the middle kymograph indicates a transcribing RNAP molecule drastically reduces its elongation speed after the leading RNAP (right) terminates its transcription and dissociates.

To determine how supercoiling contributes to the observed cooperation, we analyzed the supercoiling density and resisting torque distributions in the immediate vicinity of a transcribing RNAP molecule under the conditions of low, medium, and high initiation rates (*k*_ini_ = 0.001, 0.019, and 0.131 s^−1^) respectively (**Fig 2D**). At the low initiation rate, while the supercoiling densities upstream and downstream of a transcribing RNAP are relatively low (~ ± 0.1), they produce large positive torques around the RNAP molecule (**Fig 2D**, left column), therefore decreasing the elongation rate and frequently stalling the RNAP molecule (red bars). As the initiation rate increases, the supercoiling densities around RNAP molecules also increase, but the corresponding torque decreases and becomes negative at the highest initiation rate (**Fig 2D**, middle and right columns) due to the cancellation of opposing supercoils between adjacent RNAP molecules. Consequently, majority of RNAP molecules under these conditions move processively with high elongation rates (green bars). Therefore, the reduced resisting torque can fully account for RNAP’s cooperation in highly expressed genes. A full comparison of supercoiling density and torques under all initiation rate conditions is shown in **S3 Fig**.

To further illustrate the cooperative behavior of RNAP and its relationship with local supercoil density and torque, **Fig 2E** shows three exemplary kymographs of RNAP’s translocation trajectories at the three different initiation rates. When the initiation rate is low (0.001 *s*^−1^, **Fig 2E**, left), the overall supercoiling density of the DNA (negative upstream in blue and positive downstream in orange) is maintained at a relatively low level. However, these supercoils accumulate on both sides and generate high torques to stall the transcribing RNAP molecule frequently (vertical lines in trajectories). Interestingly, when a new RNAP molecule starts transcribing (left most trajectory at ~ 850 s in the simulation), the previously slow-moving RNAP molecule (indicated by *) started elongating faster than before and its speed became similar to that of the new RNAP molecule, indicating cooperation. Under a high initiation rate (0.068 *s*^−1^, **Fig 2E**, right), the overall supercoiling density on the DNA template is high especially in the immediate upstream and downstream regions (dark blue or orange colors), but the torque that each RNAP molecule faces is low because the difference of the supercoiling density between the two sides of an RNAP molecule is small. Note that we also observed the antagonistic behavior of RNAP in these simulations, such as the one at ~ 920 s in the middle kymograph (initiation rate = 0.009 *s*^−1^, **Fig 2E**, middle), in which when the leading RNAP molecule finishes its transcription, the speed of the trailing one reduces drastically.

To further confirm the above effect, we increased Topo I activity in the same simulations and found that it could diminish the enhancement effect as what was shown in Kim *et al*. [13] (**S6 Fig**). This result is because when Topo I activity is too strong (for example, when the Topo I binding rate is 1000 times the Gyrase binding rate), negative supercoils generated by a leading RNAP molecule will be rapidly removed even before they cancel out with positive supercoils generated by the trailing RNAP molecule.

Note that two recent theoretical models [26, 27] also reproduced the cooperation between groups of RNA polymerases. However, these two models adopted different assumptions from ours. Both studies assumed that RNAP velocity is directly dependent on torsional stress [26, 27], with one study specifying no RNAP counterrotation to allow diffusion of supercoiling [27]. These assumptions may not represent the actual biological picture, as a previous study has shown that torsional stress does not cause the measured inter-stall velocity of RNAP to vary significantly other than inducing stall [10] (see **Discussion** for details). We also showed that zero RNAP counterrotation is not necessary for a cooperative behavior: in our simulation, as long as RNAP counterrotates slowly *(k_rot_* <= 1 *s*^−1^), the cooperation remains (**S7 Fig**).

### Supercoiling accumulated in a topological domain modulates transcriptional noise

Bacterial transcription exhibits a large cell-to-cell variability [43, 44, 45, 46], allowing a population of isogenic cells to exhibit heterogenous phenotypes and responses to various stimuli [47]. The variability comes from either the stochastic nature of molecular reactions (intrinsic noise) or the variation in molecule copy numbers in different cells (extrinsic noise) [48]. Here we are interested in understanding how supercoils accumulated in a topological domain could function as an additional factor to contribute to the intrinsic noise in the mRNA production of a gene. We characterize the transcriptional noise by the Fano factor of mRNA copy number, which equals one when mRNA’s production is a stochastic, Poissonian process of random birth and death [43]. When mRNAs are produced in bursts due to random activation and inactivation as observed in both prokaryotes and eukaryotes, the Fano factor is greater than one and reflects the mRNA burst size [49].

We first simulated a 4.2 kb open DNA with a 2.4 kb gene located in the center, with the ends constrained and relaxing every 0.02 s. As shown in **Fig 3A, B, D and E**, under this condition both weak and strong promoters yield an mRNA copy number distribution with a Fano factor around 1 due to the stochastic, Poissonian transcription. When we introduce a topological barrier at 0.06 kb and 4.14 kb, which dynamically loops (for 5 min) and unloops (for 1 min) the intervening DNA (to mimic realistic chromosomal DNA dynamics), both the weak and strong promoters show a substantial drop in the mean mRNA production (**Fig 3C, F, G, H**). However, the strong promoter exhibits a larger Fano factor in mRNA copy number distribution than the weak promoter, suggesting that additional transcription noise is introduced (**Fig 3C, F**). The supercoiling density in the dynamically looping DNA exhibits strong positive or negative values, whereas that of the open DNA maintains a moderate and relatively constant level (**Fig 3G, H**, bottom panel). Notably, the mRNA production of the dynamically looping DNA is highly bursty, and each burst corresponds to the release of accumulated of torsional stress when the loop opens (**Fig 3H**, bottom panel, red shaded region).

**Fig 3.**
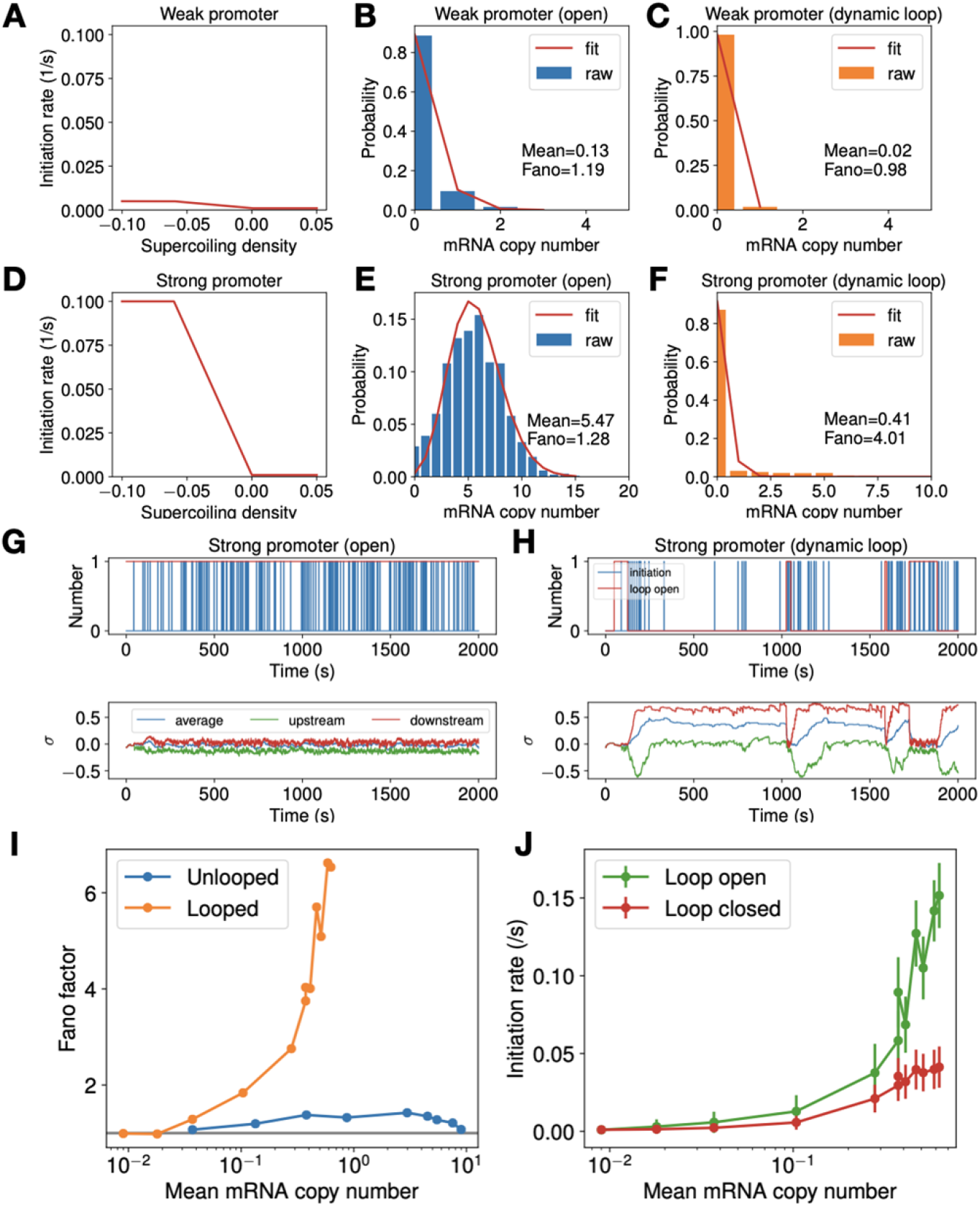
Dynamic topological domain formation results in bursty and noisy transcription from strong promoters through the accumulation and release of supercoiling. (A-F) Comparison of a weak promoter (*k_max_* = 0.005, top panel) and a strong promoter (*k_max_* = 0.1, bottom panel) in their responses to supercoil density (A and D), mRNA copy number distribution when the DNA is open (unlooped, B and E) and when the DNA dynamically loops and unloops (C and F). (G-H) Exemplary mRNA production time traces (top panel) and the corresponding supercoil density of the DNA (bottom panel) for the strong promoter in the open-DNA condition (G) and the dynamically looping condition (H). The pink-shaded regions in H indicate the time when the loop is open. (I) Comparison of the Fano factor as a function of mean mRNA copy number for genes in an open DNA (blue) and that in a dynamically looping DNA condition (orange). The grey line denotes Fano factor = 1. (J) Comparison of the average initiation rate of genes when the DNA is open (green) and when the DNA is closed (red) in the dynamically looping DNA condition.

As in all our simulations we also included the actions of Topo I and Gyrase in these simulations. However, we observed that topoisomerases cannot effectively alleviate the torsional stress generated by genes with strong promoters in a topologically constrained DNA loop. The stress is largely released only when the DNA loop opens. Consistent with this expectation, genes with strong promoters are transcribed with larger noises under the dynamically looping condition, and the Fano factor increases with the promoter strength (**Fig 3I**). Additionally, when the loop is closed, the torsional stress reduces the transcription initiation rate. In contrast, when the loop is open, the torsional stress is released and transcription initiates with high rates (**Fig 3J**). These observations further support the notion that the difference in transcription activities during the looped and unlooped states underlies bursty transcription and contributes to additional transcriptional noise in mRNA production.

We note that previous works [21, 25] attributed transcription bursting to the binding and unbinding of Gyrase in a topological domain. This mechanism is based on the assumption that Gyrase’s binding is a rare event compared to the timescale of transcription. Our model suggests that even if the binding and unbinding of Gyrase are far more frequent than transcription initiation, transcription can still be bursty due to the topological domain formation and opening. Furthermore, the timescale of domain formation and dissolution plays a significant role in bursty transcription: the more frequent the DNA switches between looped and unlooped state, the less bursty transcription is. We show that when the dwell time of looping is reduced from 5 min to 1 min, the Fano factor drops accordingly (**S8 Fig**, green to orange curve); if the DNA is always looped, the transcription will stay in a nearly dormant state as the torsional stress cannot be released effectively even in the presence of topoisomerase activities; consequently, the resulting Fano factor is down to unity (**S8 Fig**, red curve). Note that our model does not exclude the possibility of transcription bursting resulting from strong topoisomerase binding/unbinding, which is rare [50], but instead provides a mechanism that can result in transcription bursting independent of topoisomerase activities.

### Intergenic supercoiling mediates communication between two neighboring genes

Previous works show that neighboring genes could influence the transcription of each other, and the interaction is dependent on the relative orientation of the genes [15, 16, 51]. We propose that intergenic supercoiling dynamics may mediate communications between neighboring genes. To investigate this hypothesis, we simulated two genes of the same length (1.2 kb) and promoter strength (*k_max_* = 0.05 *s*^−1^) but arranged differently (convergent, divergent or codirectional) in a 14.4-kb long open DNA. We chose a length of 14.4-kb to accommodate the limited number of species allowed in Lattice Microbes. To examine the effects of supercoiling on transcriptional activities, we also simulated the cases where supercoiling is eliminated by increased topoisomerase activities (**S9 Fig**).

We first analyzed the effects of intergenic supercoiling on mRNA production with the intergenic distance held at 1.2 kb (**Fig 4A**). For convergent and divergent arrangements, since the two genes displayed the same transcription activities due to the symmetry, we only showed the mRNA distribution for the upstream gene. For codirectional genes, we examined the mRNA distribution for both the upstream (trailing) and the downstream (leading) genes. As shown in **Fig 4B**, we observed that the presence of supercoils led to a significant decrease in the mean mRNA production of the genes in the convergent arrangement (leftmost panel, orange bars) but a significant increase for genes in the divergent arrangement (middle panel, orange bars) compared to the condition when supercoils on the DNA are eliminated efficiently by topoisomerase activities (blue bars). For the upstream gene in the codirectional arrangement (second panel from the right), mRNA production only exhibited a slight decrease in the presence of supercoiling. For the downstream gene in the codirectional arrangement, however, we observed a significant decrease in mRNA production (rightmost panel). These results demonstrate that supercoiling indeed influences the transcription activity of genes in different orientations differentially.

**Fig 4.**
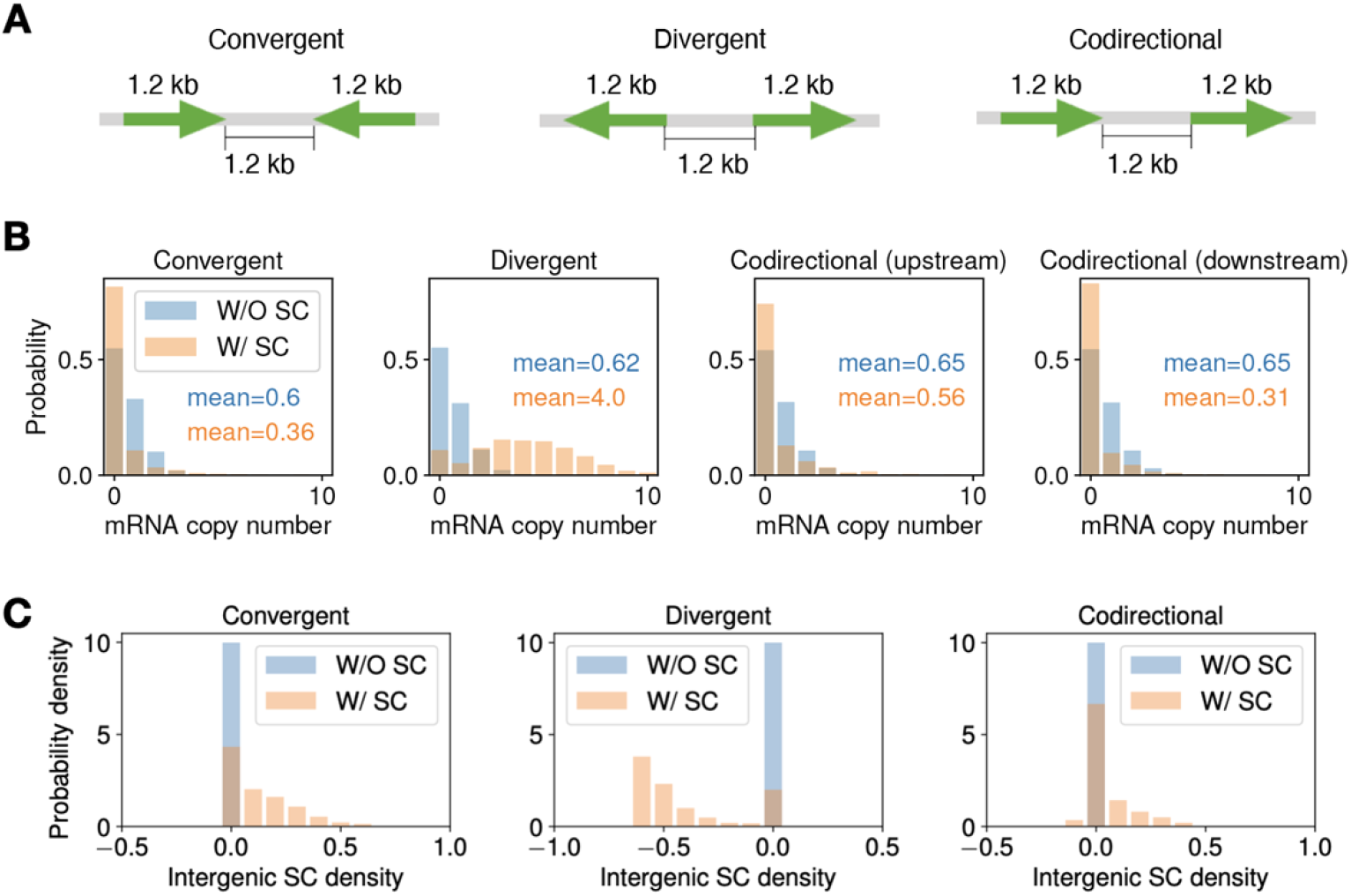
Intergenic supercoiling affects the level of mRNA production of two neighboring genes. (A) The construct for convergently, divergently and codirectionally arranged genes with the intergenic length fixed at 1.2 kb. The mRNA copy number distribution (B) and intergenic supercoiling distribution (C) for cases with (orange bar) and without (blue bar) intergenic supercoiling.

To examine how supercoiling contributes to the above observations, we plotted intergenic supercoil density distributions of these genes in **Fig 4C**. For convergent genes (left panel), the intergenic region is dominated by positive supercoils, which are produced by the transcription-induced supercoiling downstream of both genes, and hence inhibiting the transcription of both genes. In contrast, the region between divergent genes is dominated by negative supercoils (middle panel), which are produced upstream by the promoters of both genes, hence boosting transcription initiation for both genes. For codirectional genes, the intergenic region is also dominated by positive supercoiling (right panel), although the densities are lower compared to that of convergent genes. This is an interesting observation, because intuitively one would expect that positive supercoils produced by the upstream gene and the negative supercoils produced by the downstream gene would cancel each other, leading to relaxed intergenic DNA. In reality, however, the upstream gene will roughly maintain its transcription activity over time since its promoter supercoiling is frequently reset by the default negative supercoiling from the chromosome end. On the other hand, the downstream gene will experience a decline in transcription due to the immediate suppression of transcription initiation by the relaxed intergenic DNA. This difference gives rise to a positive feedback loop where the increased positive supercoiling produced by the upstream gene further decreases the initiation rates of the downstream gene. Consequently, the upstream gene does not change its mRNA production level dramatically, but the downstream gene decreases its mRNA production level.

Based on the above results, we reasoned that the presence of supercoiling should also render mRNA production of one gene sensitive to changes in the initiation rate of the other adjacent gene. To examine this hypothesis, we varied the initiation rate of one gene (**Fig 5A**) and monitored the resulting mRNA production levels of the other gene (**Fig. 5B**), the intergenic supercoiling density (**Fig. 5C**), and the correlation between the mRNA copy numbers of the two genes over time (**Fig. 5D**) for the three gene arrangements.

**Fig 5.**
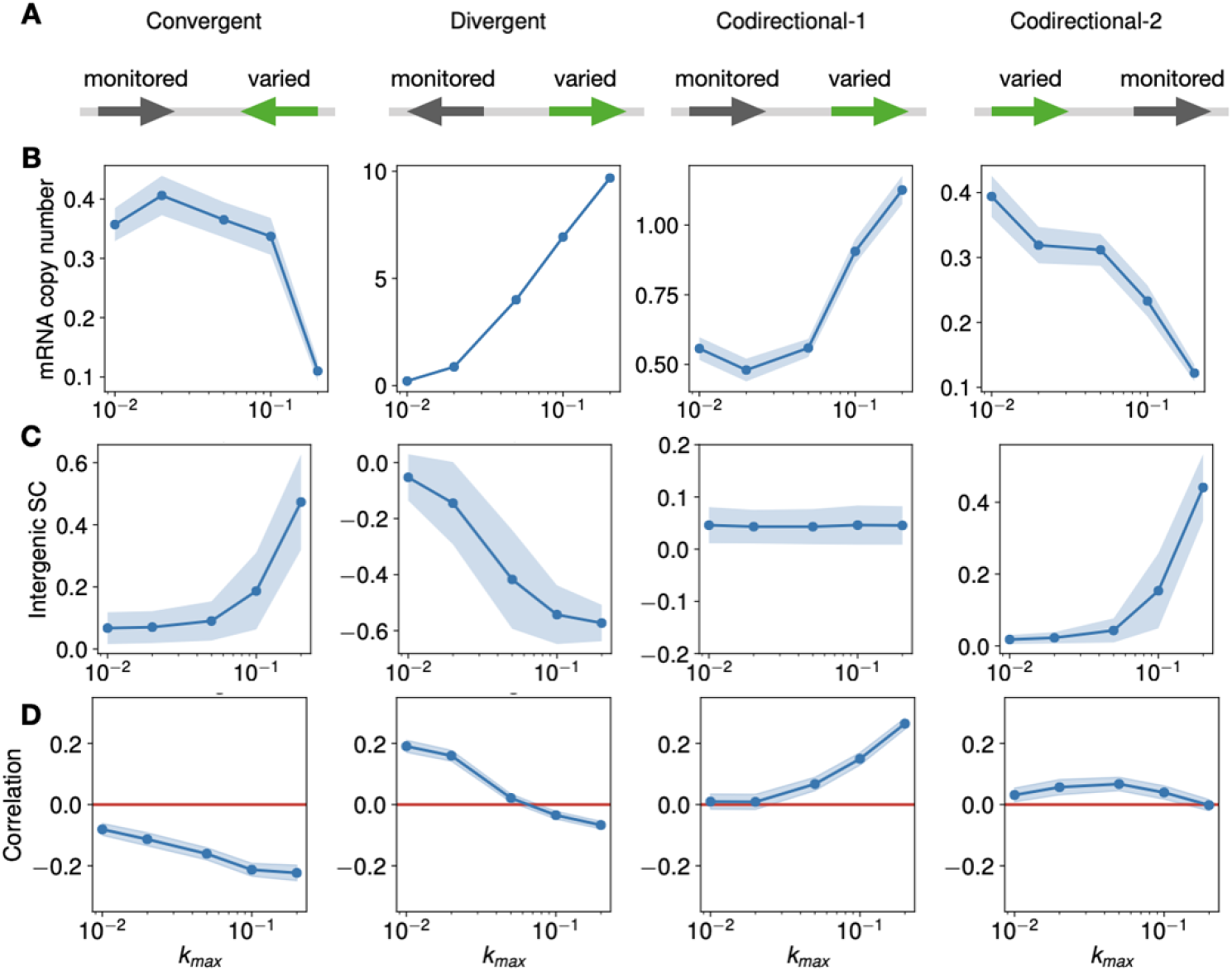
The effects of the initiation rate on expression of adjacent genes. (A) The construct for convergently, divergently and codirectionally arranged genes where the initiation rate of one gene is varied (*k_max_* = 0.01, 0.02, 0.05, 0.1, 0.2 *s*^−1^) and the initiation rate of the other gene is monitored in response to that of the first gene with an upper limit of *k*_max_ = 0.05 s^−1^ (B) The mean mRNA copy number of the monitored gene as a function of the maximum initiation rate (k_max_) of the first gene, for the four constructs shown in (A). The dot is the mean and the shaded area is mean ± SEM. (C) The intergenic supercoiling density as a function the maximum initiation rate (k_max_) of the adjacent gene. The dot is the mean and shaded area is mean ± standard deviation. (D) The Pearson correlation between mRNA copy number of two genes as a function of the maximum initiation rate (k_max_) of the gene whose initiation rate is varied. The dot is the mean and the shaded area is mean ± SEM.

For convergent genes (leftmost, **Fig 5B**), as we increased the initiation rates of one gene, the mean mRNA copy number of the other gene first remains nearly unchanged, and then decreases. This result is likely because when the initiation rate of the first gene is high (*k_max_* = 0.1 *s*^−1^ or 0.2 *s*^−1^), the intergenic region become significantly positively supercoiled (leftmost, **Fig 5C**), inhibiting the transcription of the other gene. For divergent genes (second from the left, **Fig 5B**), we observed that mRNA production increases with the initiation rate of one gene, likely due to the increased negative supercoils in the intergenic region (second from the left, **Fig 5C**). For codirectional genes, since the gene arrangement is asymmetric, we modulated the initiation rates for both the upstream gene and the downstream gene separately. When the initiation rate of the downstream gene is increased, we observed an increase in the mRNA production of the upstream gene (third from the left, **Fig 5B**), and no change in the intergenic supercoiling density (third from the left, **Fig 5C**). The homeostasis in intergenic supercoiling density suggests a feedback loop: the more the downstream gene transcribes, the more negative supercoiling is generated in the intergenic region, the more the upstream gene is activated, which produces more positive supercoils in the intergenic region to annihilate the negative supercoils. Conversely, when the initiation rate of the upstream gene is increased, we observed a decrease in the expression of the downstream gene (rightmost, **Fig 5B**), accompanied with an increased intergenic supercoiling density (rightmost, **Fig 5C**).

Lastly, to quantify the direction and strength of the interaction between two genes, we used the Pearson correlation coefficient (**Fig 5D**) between the mRNA copy number of the two genes over time. Convergent genes (left, **Fig 5D**) always maintain a negative correlation, suggesting a mutual inhibition between the two genes. In contrast, divergent genes (middle, **Fig 5D**) experience a positive correlation when the initiation rate of the adjacent gene is low to moderate (*k_max_* = 0.01 *s*^−1^ or 0.02*s*^−1^), suggesting mutual activation between the two neighboring genes. At very high initiation rate of one gene, however, the correlation between the two genes became negative, suggesting mutual inhibition between the two genes. This phenomenon is likely due to the excessive negative supercoiling in the intergenic region, which prevent RNAP to escape the promoter to engage in elongation, For codirectional genes, a positive correlation is observed when downstream gene has high transcription activity (*k_max_* = 0.1 *s*^−1^ or 0.2 *s*^−1^, third from the left, **Fig 5D**), suggesting mutual activation; in contrast, the overall correlation is very small when the upstream promoter strength is varied (right, **Fig 5D**). These results suggest a positive feedback loop between transcriptional dynamics and supercoiling mechanics in the codirectional arrangement: the transcription of the upstream gene inhibits the transcription of the downstream gene, leading to positively supercoiled intergenic region, which further inhibits the downstream transcription.

## Discussion

In this study, we modeled supercoiling-dependent transcription systems in *E. coli*. This model contains the following unique features. First, both transcription initiation and elongation are supercoiling-sensitive, while previous theoretical models only incorporated one of them [20, 21, 22, 23, 24, 25, 26, 27]. Considering that supercoiling-mediated modulation of both initiation and elongation has already been shown experimentally [8, 10, 11], incorporating both processes into the model is necessary and allows us to make accurate quantitative predictions. Second, we provided an explicit description of supercoiling-sensitive elongation by modeling the stall propensity to be torque-dependent. Ma *et al*. [10] observed that torque could regulate both RNAP’s stall propensity and the velocity for processive RNAP. However, the velocity’s dependence on torque is relatively weak (for example, the velocity only drops from about 24 *bp/s* to about 18 *bp/s* when the torque changes from −6*pN* · *nm* to 7.5 *pN* · *nm*). Hence, we argue that the elongation rate is mostly regulated by the stall propensity, and the actual change in the velocity is negligible. Third, our model is pertinent to the realistic biological context. We modeled the explicit molecular events of binding/unbinding of topoisomerases, including Topo I and Gyrase, formation/dissociation of topological domains, translocation and counterrotation of RNAP, and diffusion of supercoils along the DNA. Integrating all these biological events that occur in live cells with physiologically relevant parameters renders the model explicit in its biological implications.

Using this model, we quantitatively reproduced the orchestration between co-transcribing RNAP molecules over long distances [13]. We showed that the cooperation between RNAP molecules, manifested by the enhancement in elongation rate in highly expressed genes, is mediated by supercoiling cancellation and the consequently reduced torque generated by RNAP. We also showed that the enhancement could only be achieved when RNAP counterrotation is slow, and that TopoI is not overly active. As RNAP would face sizeable hydrodynamic drag created by nascent transcripts and associated ribosomes under physiological conditions, the counterrotation could indeed be very slow. Thus, we predict that the cooperative behavior of co-transcribing RNAP molecules could be a universal feature in *E. coli* gene transcription.

We then utilized this model to explore the role of promoter strength and the formation of topological domain on transcription regulation mediated by supercoiling. Our simulations showed that a dynamically looped topological domain could reduce the average mRNA production within the domain and add intrinsic noise to gene transcription in a promoter strength-dependent way. Thus, topological domains could act as transcription regulators and globally manipulate the distribution of transcription levels. We also provided another mechanism for transcriptional bursting by the dynamic formation and dissolution of topological domains, which is in addition to Gyrase binding/unbinding as previously proposed [21, 25]. Note that the necessary condition for supercoiling-mediated transcriptional bursting is that there is enough timescale separation in supercoiling generation and dissipation. However, the timescale Gyrase binding/unbinding is still controversial: the *in vitro* study by Chong *et al*. [8] estimated the Gyrase to bind/unbind every several minutes, while the *in vivo* experiment by Stracy *et al*. [37] observed each binding to span only 2-10 seconds. If the latter picture is true, Gyrase activity alone will not provide enough time scale separation for supercoiling generation and dissipation. On the other hand, we also lack the experimental evidence for topological domain-induced transcriptional bursting: although the stability of several transcription factor-mediated loops has been measured [52, 53, 54] (such as LacI- or CI-mediated loop, whose stability ranges from seconds to minutes), the kinetics of other kinds of topological domains (formed by nonspecific nucleoid-associated proteins like Fis and H-NS [55]) remain largely unknown. More work is needed to elucidate the mechanism underlying transcriptional bursting.

We also analyzed the modulation of mRNA production of one gene by a neighboring gene and its dependence on the relative orientation of the two genes. We showed that even a simple two-gene system could yield complex behaviors due to supercoiling dynamics (for example, the asymmetry in gene regulation in codirectional genes). These behaviors brought about the need to use supercoiling-dependent models in facilitating the design of synthetic genetic circuits. For example, for convergent genes, we notice that supercoiling brings a negative correlation to the transcription of the two genes, which means that it is possible to use convergent genes to enhance the bistability of synthetic toggle switches [56].

Finally, we point out that our model reproduced qualitatively but not quantitatively the antagonistic behavior reported by Kim *et al*. [13]. In our simulations, RNAP molecules do slow down in response to promoter inactivation when 2700 bp has been transcribed, and the torsional stress is transmitted towards downstream (left panel, **S10 Fig**). However, the slowdown effect is not as pronounced as observed in the experiments: the simulated elongation rate reduced from 33 bp/s to 26 bp/s after promoter inactivation, while the experimentally measured elongation rate reduced from 30 bp/s to lower than 10 bp/s. A sizeable slowdown could be achieved if we make the promoter inactivation earlier: if the promoter is inactivated when 300 bp is transcribed, the elongation rate of remaining RNAP molecules will reduce to about 10 bp/s (right panel, **S10 Fig**). Two possible reasons might contribute to this observation. One is that the supercoiling diffusion coefficient is too small such that the torsional stress is not transmitted fast enough to reach the most downstream RNAP before transcription termination. Another possible reason is that the binding of multiple RNAP molecules might make the DNA more rigid, as assumed in [27] -- as a result, the experienced torque by each RNAP might be larger than we modeled. Further investigations are necessary to identify and differentiate these potential causes.

## Acknowledgements

We would like to thank Dr. Taekjip Ha for useful comments on the initial draft.

